# Paralog dispensability shapes homozygous deletion patterns in tumor genomes

**DOI:** 10.1101/2022.06.20.496722

**Authors:** Barbara De Kegel, Colm J. Ryan

## Abstract

Genomic instability is a hallmark of cancer, resulting in tumor genomes having large numbers of genetic aberrations, including homozygous deletions of protein coding genes. That tumor cells remain viable in the presence of such gene loss suggests high robustness to genetic perturbation. In model organisms and cancer cell lines, paralogs have been shown to contribute substantially to genetic robustness – they are generally more dispensable for growth than singletons. Here, by analyzing copy number profiles of >10,000 tumors, we test the hypothesis that the increased dispensability of paralogs shapes tumor genome evolution. We find that genes with paralogs are more likely to be homozygously deleted and that this cannot be explained by other factors known to influence copy number variation. Furthermore, features that influence paralog dispensability in cancer cell lines correlate with paralog deletion frequency in tumors. Finally, paralogs that are broadly essential in cancer cell lines are less frequently deleted in tumors than non-essential paralogs. Overall our results suggest that homozygous deletions of paralogs are more frequently observed in tumor genomes because paralogs are more dispensable.

## Introduction

Tumor genomes typically abound with genetic aberrations, ranging from missense mutations in individual genes to deletions of entire chromosome arms (Vogelstein *et al*, 2013). These genetic alterations can result in reduced functionality, or complete loss of function, of multiple proteins in any given tumor cell. Despite these alterations, tumor cells remain viable and even thrive, suggesting that they are highly robust to genetic perturbation. This raises an important question: how do tumor cells tolerate such gene loss? One potential explanation is paralog buffering.

Paralogs are genes that arose from gene duplication events, the primary means by which new genes are created (Zhang, 2003). Many paralog pairs retain at least some degree of functional redundancy, even after long evolutionary periods, which may allow them to buffer each other’s loss (Kuzmin *et al*, 2021). In multiple model organisms, paralogs have been demonstrated to contribute significantly to genetic robustness (Gu *et al*, 2003; Kamath *et al*, 2003; White *et al*, 2013). For example, in budding yeast systematic gene deletion studies have revealed that loss of paralog genes is generally better tolerated than loss of singletons (genes without a paralog) (Gu *et al*, 2003). Subsequent double gene deletion studies revealed that ~30% of yeast paralog pairs display negative genetic interactions, where the combined gene deletion causes a greater than expected fitness defect, indicative of a buffering relationship (DeLuna *et al*, 2008; Dean *et al*, 2008; VanderSluis *et al*, 2010).

We, and others, have made similar observations using loss of function screens in human cancer cell lines – paralogs are generally more dispensable for cellular growth than singletons, and this is even more evident for highly sequence similar paralogs and genes with multiple paralogs (Dandage & Landry, 2019; De Kegel & Ryan, 2019; Wang *et al*, 2015; Dede *et al*, 2020). Furthermore many members of paralog pairs can be lost individually but not in combination, suggesting their dispensability can be directly attributed to paralog buffering (De Kegel & Ryan, 2019; De Kegel *et al*, 2021; Dede *et al*, 2020). While it is therefore clear that paralogs contribute to the genetic robustness of tumor cell lines *in vitro*, it is not clear whether this is a significant factor in the genetic robustness of tumors *in vivo*.

One means to explore the impact of paralogs on genetic robustness in tumors *in vivo* is through the analysis of compendia of tumor genomes, which provide a record of those genetic alterations that can be tolerated by tumor cells, under at least some circumstances. Tumors evolve through the accumulation of somatic alterations followed by clonal selection. Alterations that increase cellular fitness will likely be observed more frequently across tumor genomes, as positive selection would increase their prevalence in any given tumor population. Conversely, deleterious alterations will likely be observed less frequently across tumor genomes, as negative selection would cause clonal lineages with such aberrations to die out. If paralog genes are more dispensable for tumor cells *in vivo*, we expect that in general deleterious alterations of paralog genes will be under weaker negative selection than similar alterations to singleton genes. As the majority (>60%) of genes in the human genome have at least one paralog, paralog dispensability has the potential to substantially shape tumor genomes.

Analyses of selection in cancer have largely focused on positive selection – a key aim has been to distinguish driver alterations, that directly promote tumorigenesis, from passenger alterations, that provide no fitness benefit and are presumed to be present due to ‘hitchhiking’ with driver alterations (Vogelstein *et al*, 2013; Lawrence *et al*, 2014). More recently several dN/dS based approaches – which compare the expected number of nonsynonymous mutations (dN) to the expected number of synonymous mutations (dS) within a gene – have been used to identify signals of negative selection. In general, evidence for selection against missense mutations has been challenging to detect. In diploid and tetraploid regions even nonsense mutations in broadly essential genes appear to be well tolerated, but in haploid regions clear patterns of selection against nonsense mutations have been detected (Martincorena *et al*, 2017; Weghorn & Sunyaev, 2017; Van den Eynden *et al*, 2016; López *et al*, 2020), suggesting that most non-driver genes are haplo-sufficient in cancer cells.

Here we focus our analysis on homozygous gene deletions – genetic aberrations that are guaranteed to result in complete protein loss. Homozygous deletion (HD) frequency has been previously used to identify tumor suppressor genes, whose loss recurs across tumor genomes due to positive selection (Cheng *et al*, 2017; Zack *et al*, 2013); but here we are particularly interested in HD frequency of non-driver (i.e. passenger) genes, whose loss will likely not provide a selective advantage. In the absence of positive selection recurrent HDs may still be observed due to localized decreased negative selection strength and/or increased HD generation rate, which can occur at fragile sites or near telomeres (Bignell *et al*, 2010; Cheng *et al*, 2017). If paralog HDs are subject to weaker negative selection, this should be observable as a higher frequency of HDs among paralog compared to singleton genes – assuming equal rates of HD generation. In other words, we expect that tumor clones with stochastically acquired HDs of singleton genes will expire at a higher rate than clones that acquired HDs of paralog genes, leading to different observed frequencies of these types of HDs across tumor genomes (Fig. 1).

**Fig 1.**
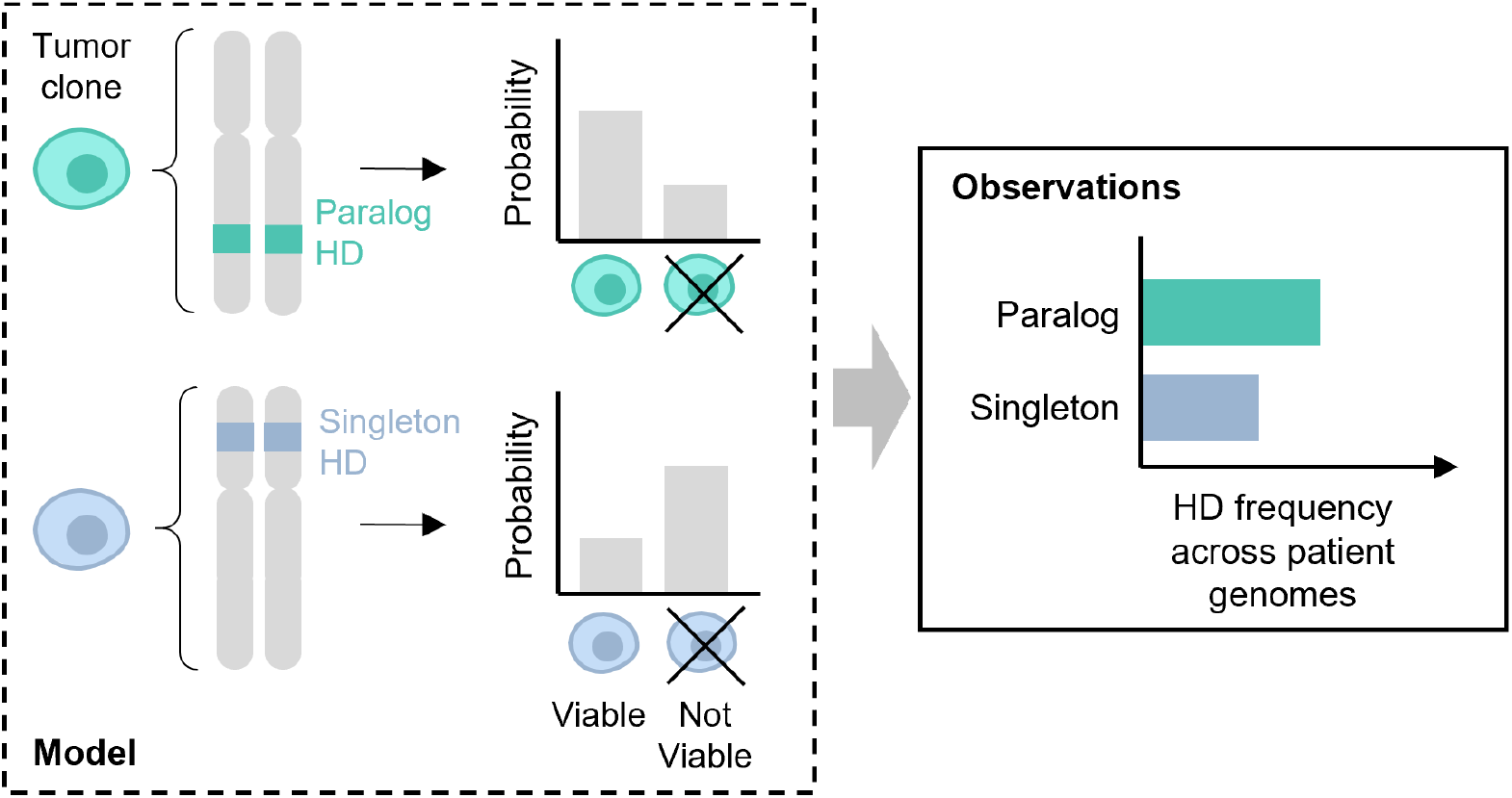
Theoretical model for the observed HD frequencies of paralog and singleton passenger genes. Tumor clones with homozygous deletion of a paralog gene are more likely to be viable than tumor clones with homozygous deletion of a singleton gene. This leads to a comparatively higher frequency of paralog HDs across patient samples.

To investigate patterns of HDs we perform a systematic analysis of homozygous deletions in 9,966 tumor samples from The Cancer Genome Atlas (Cancer Genome Atlas Research Network *et al*, 2013) and 1,782 tumor samples from the International Cancer Genome Consortium (ICGC/TCGA Pan-Cancer Analysis of Whole Genomes Consortium, 2020). We show that among non-driver genes, HDs are more likely to be observed in paralogs than singletons and that this is not solely due to factors known to influence HD generation, such as proximity to recurrently deleted tumor suppressors or fragile sites. Furthermore, we find that properties of paralogs previously shown to influence their dispensability in cancer cell lines also influence their homozygous deletion frequency in tumors, e.g. HDs are more likely to be observed in genes that have a larger number of paralogs. Finally, we show that HDs are less likely to be observed for essential paralogs than non-essential paralogs, suggesting that the increased HD frequency of paralogs is due to their generally higher dispensability rather than being a general feature of paralogs.

In addition to supporting the hypothesis that paralogs are a significant source of robustness in tumors, our results provide insight into the homozygous deletion patterns observed in tumors and highlight constraints operating on gene loss in tumor cells.

## Results

### Homozygous deletions are more likely to be observed for paralog than singleton passenger genes

To assess the homozygous deletion frequency of paralog and singleton genes, we first obtained allele-specific copy number profiles for 9,966 tumor samples spanning 33 cancer types from the Cancer Genome Atlas (TCGA) (see Methods). Among these samples less than half (4,252) had at least one autosomal homozygous deletion (HD) segment – defined as a segment of the genome where the copy number of both alleles is zero – and fewer again had a HD of a protein-coding gene. To ensure that only complete protein loss was considered, we conservatively called protein-coding genes as deleted when the full coding sequence of their longest associated transcript was deleted. After removing a small number of ‘hyper-deleted’ samples (see Methods) we found that 2,780 samples contained an HD of at least one autosomal protein coding gene (Fig. S1A, Table S1).

We visualized gene HD frequency across the whole genome and observed that, consistent with expectation, several well-known tumor suppressor genes (TSGs), including *CDKN2A, PTEN, RB1* and *SMAD4* are associated with peaks of recurrent HDs (Fig. S2). To identify these genes systematically we assembled a list of 652 known cancer driver genes (TSGs and oncogenes) by combining the Cancer Gene Census (Sondka *et al*, 2018) with a comprehensive set of driver genes identified in the TCGA (Bailey *et al*, 2018) (see Methods). We found that roughly half, or 188, of the TSGs are fully homozygously deleted at least once in the TCGA samples, while 77 are recurrently homozygously deleted (in three or more samples). As our primary interest is not in driver genes, and as they are likely subject to different selection pressures than other genes, we excluded them from our analysis. The exclusion of all 652 driver genes, plus 16 genes whose coding regions overlap the coding regions of these driver genes, left us with 16,898 genes, that we refer to as passengers, for further analysis (see Methods).

We find that most of these genes are never homozygously deleted (Fig. 2A, top). Specifically, ~66% of all passenger genes have no HDs in any TCGA tumor samples, while only ~9% of passenger genes are recurrently deleted – defined as having HDs in at least three samples. We note that due to the limited sample size, this is likely an underestimate of the number of passenger genes that can be deleted in tumors. To estimate the level of saturation we sub-sample the TCGA dataset and plot the number of unique genes with at least one observed HD for increasing sample sizes (Fig. S3). We observe that up to ~3,000 samples the number of unique gene HDs increases linearly with sample size and thereafter the rate of newly seen gene HDs starts to decrease, with one new gene HD being observed for every 2-3 samples added to the cohort. However, we are not close to saturation, meaning that it is highly likely that some passenger genes are never homozygously deleted due to chance rather than negative selection.

**Fig 2.**
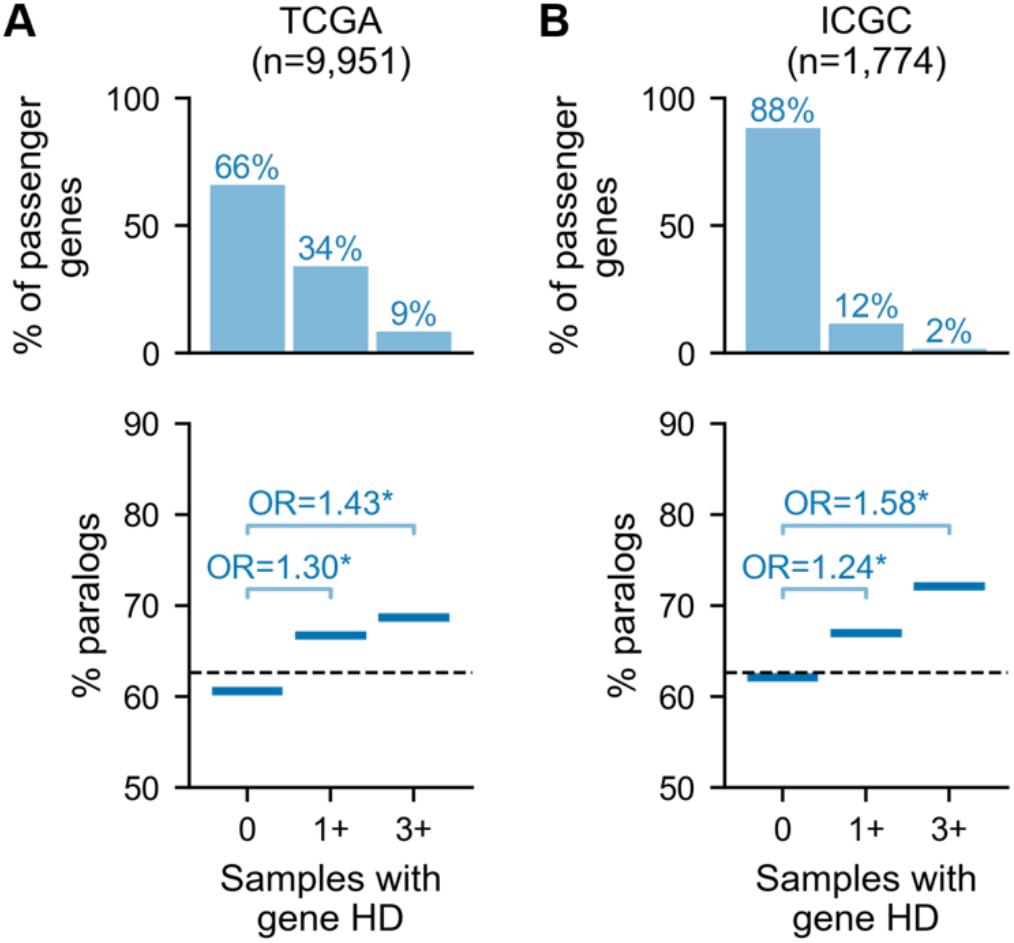
Homozygously deleted passenger genes are enriched in paralogs. (A) Top: Percentage of passenger (non-driver) genes that are deleted in 0, at least 1 and at least 3 TCGA tumor samples. The number of TCGA tumor samples considered is shown. Bottom: For passenger genes grouped by their number of HDs, the percentage of genes in each group that are paralogs (note: the 3+ group is a subset of the 1+ group). Annotations show the Odds Ratio (OR) for a Fisher’s Exact Test comparing the percentage of paralogs among genes with 0 vs. 1+ HDs and 0 vs. 3+ HDs; asterisk (*) indicates *p*<0.05. The dashed line shows the percentage of all passenger genes that are paralogs. (B) Same as (A) but for ICGC tumor samples.

Genome-wide it is clear that gene HD frequency is generally low, with a small number of highly deleted regions corresponding to TSG locations (Fig. S2). This paucity of HDs is in line with a previous analysis of a different set of tumor samples which showed that compared to expectation (based on hemizygous deletion frequency), HDs are generally strongly depleted across the genome, likely due to negative selection (Cheng *et al*, 2017). As the frequency of passenger gene HD is low (median zero HDs per passenger gene) and highly skewed, we focus our analysis on the odds of observing any HD and recurrent HD.

Grouping passenger genes according to the number of samples in which they are homozygously deleted, we find that, compared to singletons, paralogs are significantly enriched among passenger genes that are deleted at least once (Fisher’s Exact Test (FET): Odds Ratio=1.30, *p*=4e-15) and further enriched among recurrently deleted genes (FET: OR=1.43, *p*=2e-9; Fig. 2A).

We validate this finding in an independent cohort of 1,782 tumor samples from the International Cancer Genome Consortium (ICGC) (Fig. 2B, Table S2; see Methods). As this cohort is smaller, fewer passenger genes (~12%) are observed to be homozygously deleted at least once (Fig. 2B, top) – nevertheless, we again observe that paralogs are enriched, with increasing magnitude, among ever (FET: OR=1.24, *p*=2e-5) and recurrently deleted passengers (FET: OR=1.58, *p*=7e-4; Fig. 2B). To assess whether this trend is also evident within tumors from individual cancer types, we separately analyzed each TCGA cancer type with at least 600 samples (n=9). As the overall number of HDs is lower with the smaller sample sizes, we only compared genes that are never deleted vs. genes with at least one deletion in each sample subset. For 8 out of 9 cancer types we observed that paralogs are significantly enriched among passengers with at least one HD (Fig. S4A). This observation indicates that the increased HD frequency of paralogs compared to singletons is robust to cancer type specific copy number alteration biases, and to overall HD burden, which can vary considerably by cancer type (Cheng *et al*, 2017; Zack *et al*, 2013). For one cancer type (Colon Adenocarcinoma, COAD) the trend appears consistent, but the overall frequency of HDs is lower, and the enrichment is not statistically significant.

We next asked whether paralogs are also subject to more hemizygous deletions than singletons, which could lead to unequal rates of HD generation. To assess this we identified all genomic segments with loss-of-heterozygosity (LOH), i.e. all segments where one (but not both) of the alleles has copy number 0. We do not find that paralogs are more frequently subject to LOH than singletons in either the TCGA or ICGC cohort (Fig. S4B-C); when considering all LOH segments we even see that singletons are slightly more frequently subject to LOH in the ICGC cohort (Fig. S4C, left), but when considering only focal LOH segments – i.e. segments whose length is less than half of the chromosome arm’s length, which is the case for all HD segments – there is no significant difference between paralog and singleton LOH frequency in either cohort. In strong contrast to what we saw for homozygous deletions, all passenger genes are hemizygously deleted in at least one tumor sample (in both cohorts), and most are frequently hemizygously deleted, underscoring that hemizygous gene loss appears to be well-tolerated (Martincorena *et al*, 2017; Van den Eynden *et al*, 2016). This broadly suggests that paralogs are not in general more prone to deletions than singletons – we investigate this further in the next section.

### Paralogs are still more frequently homozygously deleted after accounting for proximity to tumor suppressors and fragile sites

Having established that HDs are more frequently observed for paralogs than singletons, we next wished to understand whether this could be attributed to other genome features, as copy number alterations of specific genomic regions can be influenced by multiple factors. In tumors perhaps the most obvious influence is the presence of tumor suppressor genes (TSGs) whose loss may be subject to positive selection, driving up local deletion frequency (Cheng *et al*, 2017; Zack *et al*, 2013). However, there are additional factors that are known to influence copy number variation, including proximity to fragile sites, telomeres, and centromeres. Visual analysis of the observed gene HD frequency across the genome (Fig. S2) suggests that these factors are in some cases associated with higher HD frequency and thus they may explain the relatively higher HD frequency of paralogs – e.g. perhaps paralogs are simply enriched in regions close to fragile sites. Here we discuss each of the factors in turn, and then account for them collectively in our analysis.

Firstly, paralogs may be more frequently homozygously deleted if they are in general located closer to recurrently deleted TSGs (Fig. 3A, S2). Typically an HD results in the loss of several chromosomally adjacent genes – in the TCGA dataset a median of three genes are lost per gene-deleting HD segment. Genes adjacent to TSGs might therefore be deleted more frequently due to positive selection for the loss of the TSG. For example, multiple passenger genes adjacent to the recurrently deleted tumor suppressor *TGFBR2* are themselves also recurrently deleted, including *ZCWPW2* and *GADL1* (Fig. 3A).

**Fig 3.**
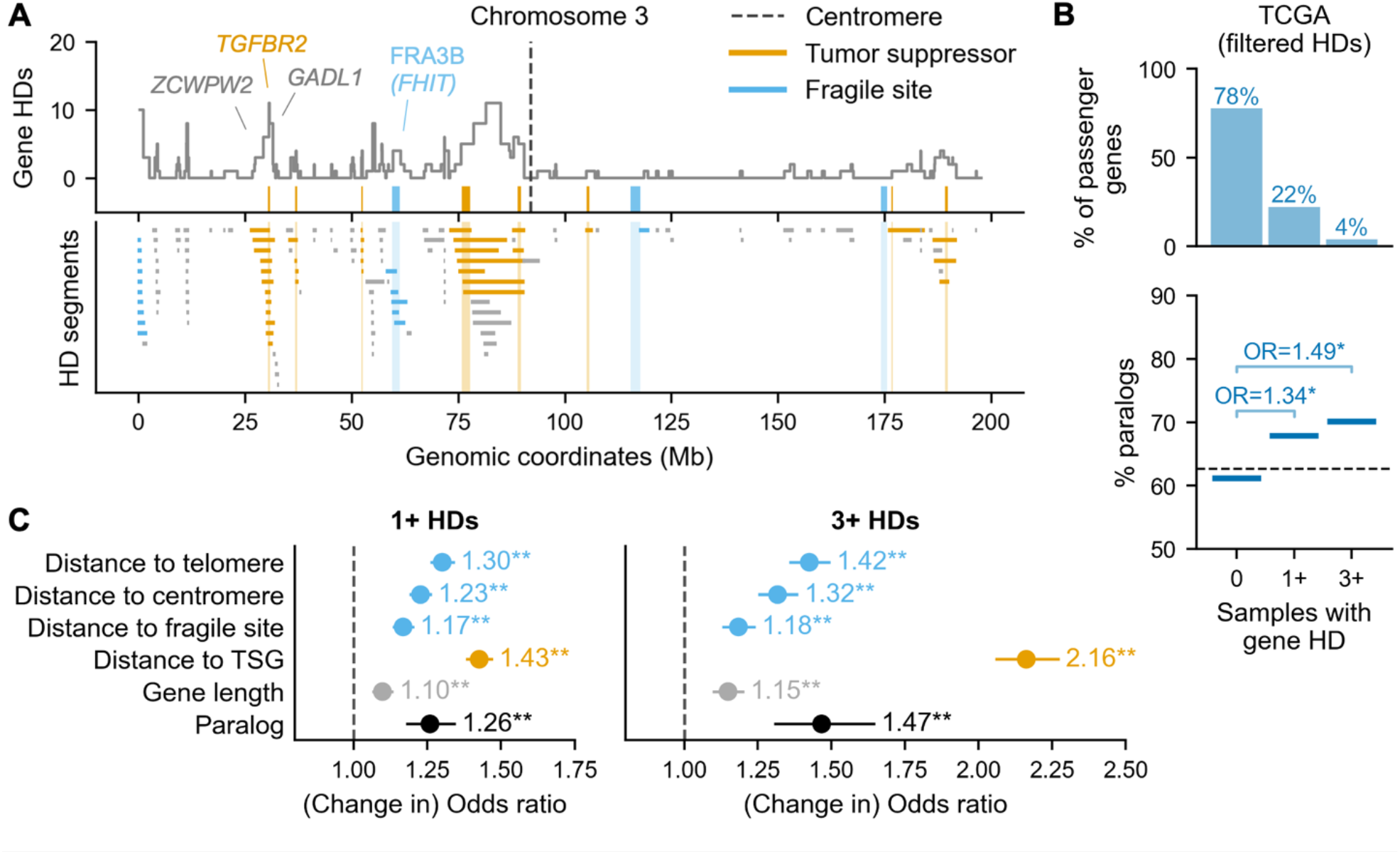
Having a paralog independently increases the odds of observing 1+ HDs for a passenger gene. (A) Top: Line plot showing the number of TCGA samples in which each gene along chromosome 3 is homozygously deleted. Orange ticks show the location and width of TSGs that are deleted in at least 1 sample. Blue ticks show the location and width of fragile sites. Bottom: Location of all HD segments affecting chromosome 3. Segments are colored according to whether they overlap a TSG (orange) or a fragile site / telomere region (blue). (B) Same as Fig. 2A but calculated after excluding all segments that overlap a TSG, fragile site, telomere or centromere, i.e., the colored segments in (A). (C) Odds ratio estimates for the association of paralogy and other genomic factors with observing 1+ HDs (left) and 3+ HDs (right) vs. 0 HDs of a passenger gene across tumor samples from TCGA and ICGC combined. For all variables except paralogy the dot represents the change in odds ratio for a one standard deviation increase in the variable value. Lines indicate 95% confidence intervals and asterisks indicate *p*-values from the logistic regression (** = *p*<0.01, * = *p*<0.05).

A second factor that could influence gene deletion frequency is proximity to fragile sites – chromosomal regions that are particularly prone to instability (Glover *et al*, 2017). Previous work has shown that some of the deletion hotspots in cancer genomes can be attributed to fragile sites (Cheng *et al*, 2017; Zack *et al*, 2013; Bignell *et al*, 2010). Genes close to fragile sites may thus be more likely to be deleted at least once in the evolutionary history of each tumor. To assess the impact of fragile sites we obtained genomic coordinates for 15 major autosomal fragile sites identified in the PCAWG study (Li *et al*, 2020) – these sites were identified based on analyses of (a subset of) the TCGA and ICGC tumor samples used in this work. The fragile sites all overlap large (>600kb) genes, some of which, such as *FHIT* (Fig. 3A), have also been identified as tumor suppressors – as was noted in (Glover *et al*, 2017). We observe that, although HDs from the TCGA cohort do recur with relatively high frequency at these fragile sites, they are much shorter than other HD segments (mean length ~0.34Mb vs. ~0.83Mb) and are mainly intragenic, i.e. they rarely result in even one full gene HD (Fig. S5A). However, given the low frequency of HDs overall, even small HD peaks, which can be observed over some fragile sites (Fig 3A, S2), are important to take into account.

The third factor to consider is proximity to telomeres and centromeres, as (sub)telomeric and centromeric regions have been shown to be enriched for deletions both in cancer genomes specifically (Zack *et al*, 2013; Li *et al*, 2020; Cheng *et al*, 2017; Beroukhim *et al*, 2010), and across human genomes in general (Collins *et al*, 2020). As with fragile sites, HD generation could thus be higher for genes close to these regions – indeed, for several chromosomes we observe gene HD peaks at the chromosome edges or centromeres that are not explained by a known TSG or fragile site, e.g. at the start of chromosome 3 (Fig 3A, Fig S2).

To account for the influence of the potentially confounding factors described above, we first repeated our analysis of passenger HDs (from Fig. 2A) after excluding all HD segments that at least partially delete a TSG, at least partially overlap a fragile site, or are telomere- or centromere-bound (see Methods). Fig. 3A highlights the HD segments that are dropped among all HD segments on chromosome 3 that fully delete at least one passenger gene. In total the excluded segments make up ~55% of all passenger-deleting HD segments in the TCGA data, which is mainly attributable to TSG associated segments. As expected, when considering just the remaining HDs, the percentage of passenger genes with no HDs is higher, while the percentage of recurrently deleted passenger genes is lower (Fig. 3B, top). Nevertheless, we find that paralogs are enriched among ever deleted passenger genes and even more enriched among recurrently deleted passenger genes (Fig. 3B).

As fragile sites appear to be centered on large genes, and some additional recurrently deleted regions have been identified over large genes in tumor samples (Beroukhim *et al*, 2010; Glover *et al*, 2017), we considered that gene length (i.e. number of bases from the start of the first to end of the last exon for the longest transcript) might also be a potential confounder – particularly because we, and others (Ibn-Salem *et al*, 2017), observe that paralogs are on average significantly longer than singletons (mean gene length ~58 kb vs. ~39 kb, Mann-Whitney U (MWU) test, *p*<1e-16). However, as we only count full gene HDs, the impact of gene length on the likelihood of observing a gene HD is likely limited – the correlation between gene length and HD frequency is only 0.03 (Spearman’s correlation coefficient, *p*=6.5e-5), compared to 0.16 (*p*<1e-16) when counting all partial gene HDs.

To understand the relative contribution of paralogy and the other identified factors to HD frequency, we fit logistic regression models for ever and recurrent gene deletion (i.e. 0 vs. 1+ and 0 vs. 3+ HDs) that integrate paralogy, gene length (which we could not account for in Fig. 3B) and distance to the nearest fragile site, telomere, centromere, and recurrently deleted TSG (Fig. 3C, Table S3; see Methods). The four distance variables are inverted so that higher values indicate the gene is closer to the region of interest and capped at 10Mb – this corresponds to the ~99th percentile of all observed gene-deleting HD segment lengths and thus represents a plausible range of influence (Fig. S5B, see Methods). To increase statistical power for this regression analysis we combine the TCGA and ICGC cohorts into one dataset by summing gene HDs. For the “distance to recurrently deleted TSG” variable we thus consider distance to 90 TSGs that have at least three HDs across this joint dataset.

We find that paralogy is independently associated with higher odds of observing any HD for a passenger gene (OR=1.26, Fig. 3C), and that the increase in deletion odds associated with paralogy is greater for recurrent HDs (OR=1.47) – corroborating what we observed earlier. For both the ever and recurrent HD models, the full model describes the data significantly better than a model that omits paralogy (Likelihood ratio tests, *p*<1e-8). As might be expected, proximity to the nearest recurrently deleted TSG is the single most influential factor – a one standard deviation increase in this variable (which corresponds to ~1Mb) increases the odds of observing any HD by a factor of 1.43 and the odds of observing recurrent HD by a factor of 2.16. The odds associated with the other genomic factors are similar to or smaller than those associated with paralogy.

Overall the logistic regression models for any and recurrent passenger gene HD suggest that having at least one paralog significantly increases the likelihood of observing at least one HD for a given gene, independently of the influence of positively selected TSG HDs, gene length or proximity to genomic fragile sites (including centromeres and telomeres). Being a paralog increases the odds of an HD ever being observed in the cohort by a factor of 1.26 and the odds of a recurrent HD in the cohort by a factor of 1.47.

### Homozygous deletion frequency of paralog passengers is influenced by paralog properties

In previous work we and others observed that certain features of paralog genes make them more or less likely to be dispensable for the growth of cancer cell lines (De Kegel & Ryan, 2019; Dandage & Landry, 2019). Specifically, we found that genes from larger paralog families and genes with a more sequence similar paralog are more likely to be dispensable – presumably because those properties can be linked to increased buffering capacity for paralog loss. We also found that paralogs that originated from a whole genome duplication (WGD) as opposed to a small scale duplication (SSD) were both more likely to be essential at least some of the time, but less likely to be essential all of the time – this could be explained by WGD paralogs being more likely to be synthetic lethal (De Kegel *et al*, 2021; De Kegel & Ryan, 2019). Having determined that paralog passenger genes are overall more likely to be homozygously deleted in tumors, we next asked whether there are differences in the deletion frequency of paralogs with different properties.

To answer this question we first compared family size for paralog passenger genes grouped according to whether they are ever and/or recurrently deleted. We find that paralog genes with at least one HD across tumor samples from both cohorts are more likely to come from a big paralog family, i.e. have more than the median number of paralogs (3), than those that are never deleted (FET: OR=1.25, *p*=5e-8; Fig. 4A). Furthermore, the odds of belonging to a big family increase for recurrently deleted paralogs (FET: OR=1.42, *p*=6e-8). This is in line with what was previously observed for paralog dispensability in cancer cell lines.

**Fig 4.**
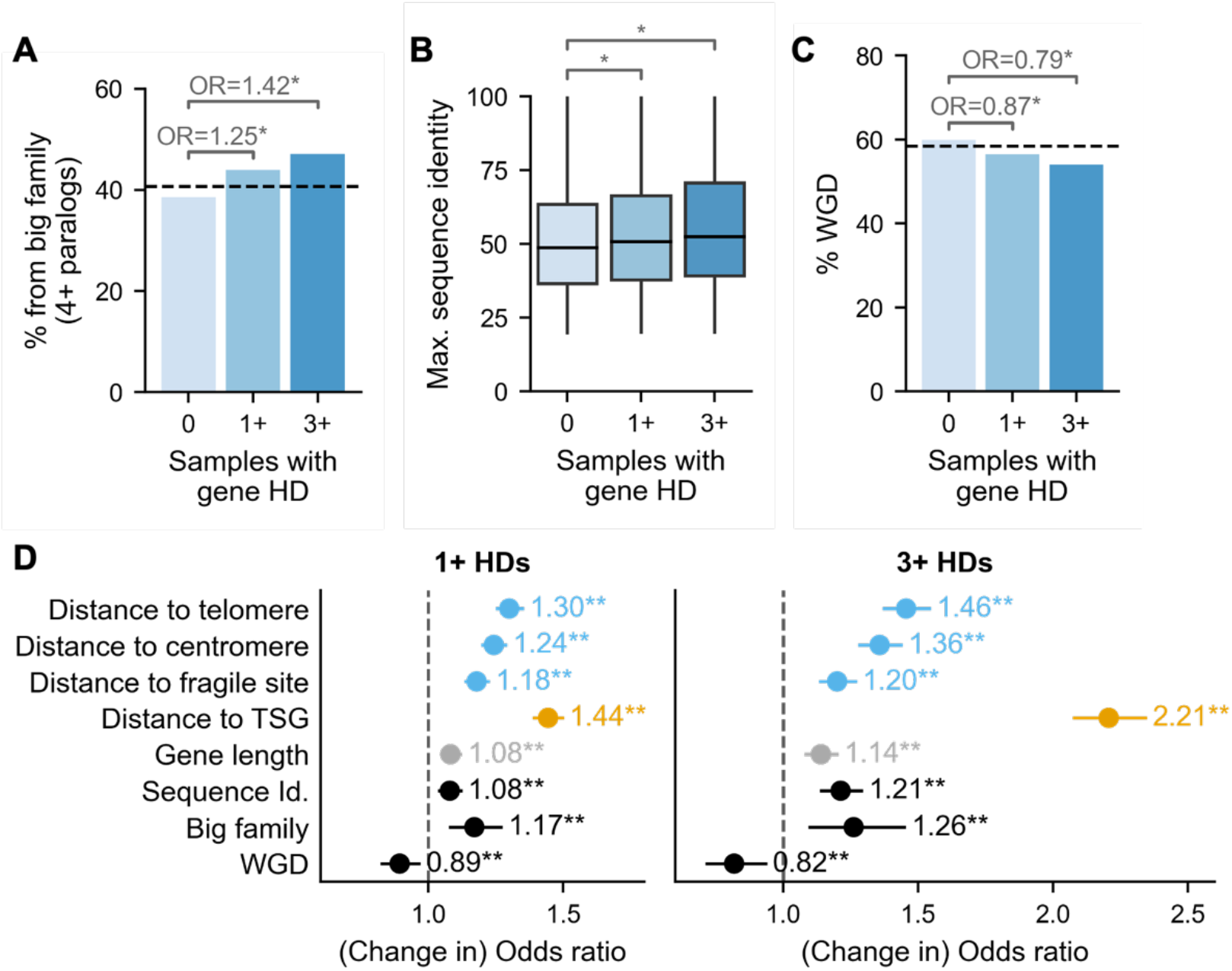
Paralog properties correlate with HD frequency. (A) Bar plot showing the percentage of paralog genes that are part of a big paralog family (i.e. have at least 4 paralogs) among paralog passengers with 0, 1+ or 3+ HDs across TCGA and ICGC samples. Annotations show the Odds Ratio (OR) from FETs comparing paralogs with 0 vs. 1+ HDs and 0 vs. 3+ HDs; asterisk (*) indicates *p*<0.05. (B) Box plots showing the maximum sequence identity, i.e. sequence identity with the closest paralog, for paralog passengers with 0, 1+ or 3+ HDs; asterisk (*) indicates *p*<0.05, MWU test. The boxes represent the first and third quartiles (Q1 and Q3) of the distribution, the horizontal black line the median, and the whiskers extend up to 1.5 * the interquartile range past Q1 and Q3. (C) Similar to (A): bar plot showing the percentage of whole genome duplicates (WGDs) among paralog passenger genes with 0, 1+ or 3+ HDs. (D) Similar to Fig. 3C: Odds ratio estimates for the association of sequence identity, big paralog family, WGD and other genomic features with observing 1+ HDs (left) and 3+ HDs (right) vs. 0 HDs of a paralog passenger gene across tumor samples from TCGA and ICGC.

We next compared, for the same groups of paralog passengers, the sequence identity each gene shares with its closest (i.e. most sequence identical) paralog (Fig. 4B). We observe that, on average, sequence identity is slightly higher for passengers with at least one HD compared to those with no HDs (mean ~52% vs. ~54%, MWU test: *p*=2e-6), and higher again for passengers with recurrent HD (mean ~56%, MWU test for 0 vs. 3+ HDs: *p*=3e-9). This is again consistent with what we observed in cancer cell lines and suggests that passenger genes with a potentially more functionally similar paralog are more likely to be dispensable.

Thirdly we asked whether we could observe differences in the HD frequency of WGD vs. SSD paralogs – we annotate individual paralog genes as WGD if they are part of at least one WGD pair, and as SSD otherwise. As WGD paralogs are more likely to be conditionally essential than SSD paralogs there is no clear expectation, based on what we saw in cancer cell lines, for whether the loss of an individual WGD gene will be more or less dispensable. We find that passengers with at least one HD are depleted in WGDs compared to paralogs with no HDs (FET: OR=0.87, *p*=0.0005) and that passengers with recurrent HD are further depleted in WGDs (FET: 0 vs. 3+ HDs: OR=0.79, *p*=0.0002, Fig. 4C); thus WGD paralogs appear to be less dispensable for tumors than SSD paralogs.

To assess whether family size, sequence identity and duplication mode (WGD vs. SSD) independently influence the probability of observing a passenger gene HD – given that these properties are to some extent correlated – we fitted similar logistic regression models to the ones described in the previous section. However, this time, as we are interested in understanding variation among paralogs, we restricted our analysis to paralog genes (i.e. excluded singletons) and added terms for each of the three paralog properties (see Methods). We find that, to varying extents, each paralog property significantly affects the odds of observing any HD, and that the magnitude of the odds ratios associated with each property increases when considering recurrent HD (Fig. 4D). Considering the 1+ HD model we find that the full model fits the data significantly better than models which omit one of the three paralog properties (Likelihood ratio tests for full model vs. model without: WGD, *p*=0.008; big family, *p*=0.0003; sequence identity, *p*=0.0004) – suggesting that each paralog property has a significant independent influence on paralog HD frequency.

### Essential paralogs are less frequently homozygously deleted than non-essential paralogs

While paralogs are in general more dispensable, certain paralogs are instead essential for cellular growth. If the key factor contributing to the increased frequency of paralog vs. singleton HDs is paralog dispensability, we would expect to see that essential paralogs are deleted less frequently than non-essential paralogs. Such an observation would indicate that higher homozygous deletion frequency is not a blanket property of paralog genes, but rather tied to their dispensability.

To estimate gene essentiality, we first use a list of broadly essential genes identified from genome-wide CRISPR screens in 769 cancer cell lines (De Kegel *et al*, 2021). These are genes that are associated with a substantial reduction in cellular growth in at least 90% of screened cell lines (see Methods). Grouping paralog passenger genes according to whether they are ever and/or recurrently deleted in the TCGA cohort (similar to Fig. 2A, but considering paralog genes only), we find that essential paralogs are significantly depleted among paralog passengers with any HD (Fig. 5A, top; OR=0.42, *p*=6e-7), and additionally depleted among paralog passengers with recurrent HD (OR=0.14, *p*=3e-6). Thus, while HDs are more likely to be observed across tumor samples for paralogs as a whole, they are less likely to be observed for cell-essential paralogs.

**Fig 5.**
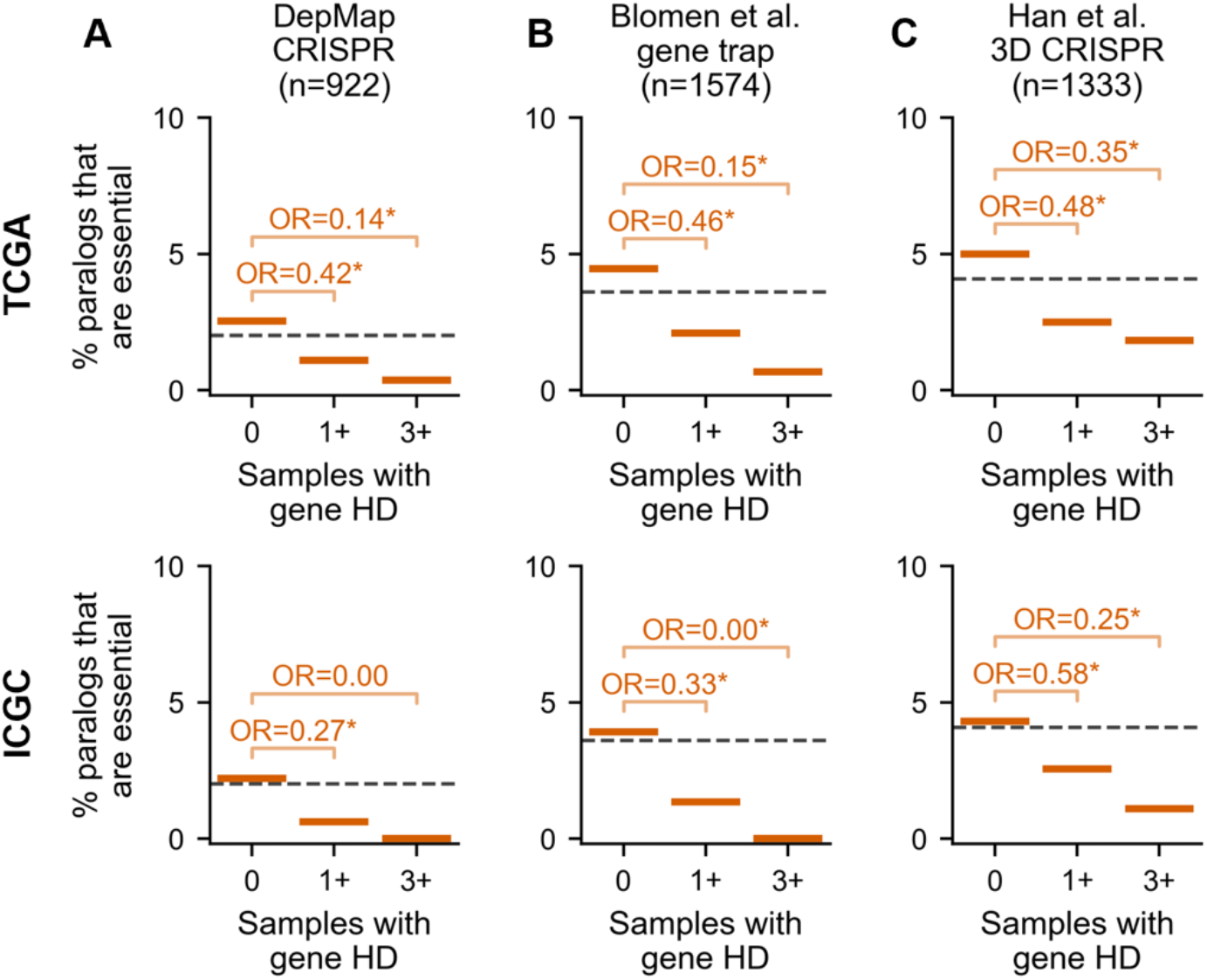
Essential paralog passengers are less likely to be homozygously deleted than non-essential paralog passengers. (A) The percentage of paralog genes that are essential, according to dependency data from DepMap CRISPR screens, among paralog passengers with 0, 1+ or 3+ HDs in the TCGA (top) or ICGC (bottom) cohort. The total number of essential paralog passengers identified from the DepMap CRISPR screens is shown; the percentages are calculated in reference to all paralog passengers for which DepMap essentiality data was available. (B) Same as (A) but for essentiality data from Blomen *et al*. gene trap screens (C) Same as (A) but for essentiality data from Han *et al*. 3D CRISPR screens.

We confirmed this tendency using two other sets of essential genes: the 1,734 genes identified as essential from gene trap screens performed in KMB7 and HAP1 cell lines (Blomen *et al*, 2015) (Fig. 5B, top); and the overlap of the top 2000 negative hits from genome-wide CRISPR screens in three lung cancer cell lines grown in 3D spheroids (Han *et al*, 2020) (Fig. 5C, top; see Methods). Although growth phenotypes in vitro, even in 3D, do not perfectly recapitulate such phenotypes in vivo, the consistently strong pattern for each of the three essentiality datasets suggests that they provide a reasonable approximation. We again validated our findings with tumor samples from the ICGC cohort (Fig. 5, bottom). Overall this suggests that the key reason for the increased HD frequency of paralogs vs. singletons is paralog dispensability.

## Discussion

In this work we found that, across large patient cohorts, homozygous deletions are more likely to be observed for paralog than singleton non-driver genes. The influence of paralogy on HD frequency is independent of factors that may increase genomic copy number, such as proximity to frequently deleted TSGs or fragile sites. In addition, we found that properties of paralogs that are associated with increased dispensability in cancer cell lines are also associated with increased HD frequency across tumor genomes. Finally we found that HDs are less likely to be observed for essential compared to non-essential paralogs – thus, rather than being a blanket property of paralog genes, increased HD frequency appears to be tied to gene essentiality. Overall we conclude that HDs are more likely to be observed for paralogs because paralogs are on average more dispensable and their loss is thus not selected against as strongly as singleton loss.

Most previous studies of negative selection used some form of normalized dN/dS ratio to quantify selection strength and found that, in contrast to positive selection, the signal of negative selection appears to be largely absent, outside of inactivating mutations in essential genes in haploid regions (Martincorena *et al*, 2017; Weghorn & Sunyaev, 2017; Bakhoum & Landau, 2017; Van den Eynden *et al*, 2016; López *et al*, 2020). One potential reason for this is that, given current sample sizes, there is lower statistical power to detect a significant depletion of aberrations in comparison to a significant enrichment (Zapata *et al*, 2018; Weghorn & Sunyaev, 2017). To address this limitation, we focused on HD frequency as a blunt tool for assessing the relative strength of negative selection acting on gene loss. We anticipated that negative selection might be more apparent for HDs, which result in complete protein loss, than for non-synonymous mutations, particularly as many genes essential for cellular growth appear to be haplo-sufficient (Van den Eynden *et al*, 2016; Wang *et al*, 2015). While our approach does not allow us to pinpoint specific cases where gene loss is selected against, we could show that in general deletion of paralogs appears to be under weaker negative selective pressure.

The impact of negative selection could also appear to be limited if weakly deleterious mutations are not weeded out – this can occur when, due to the lack of recombination during (asexual) tumor cell reproduction, deleterious mutations are co-inherited with alterations that confer a strong fitness advantage (Tilk *et al*, 2019). We observed that proximity to a frequently deleted TSG has a strong impact on HD likelihood for non-driver genes (Fig. 3C), but it is unclear whether this excludes influence from negative selection. It is possible that the observed TSG-targeting HDs were retained in part due to the dispensability of the co-deleted genes (McFarland *et al*, 2014). In support of this hypothesis, Pertesi et al. showed that, for several TSGs, the frequency of passenger HDs decreases approximately linearly between the tumor suppressor and the nearest essential genes on either side (Pertesi *et al*, 2019).

As paralogs make up over 60% of protein-coding genes, we propose that their increased dispensability could also in part explain the low level of negative selection that has been previously observed using dN/dS approaches (Martincorena *et al*, 2017; Weghorn & Sunyaev, 2017). However, whether the increased dispensability of paralogs in tumors can be attributed directly to paralog buffering relationships remains to be determined. In general, genes can be lost either because of robustness, of which paralog redundancy is one mechanism, or because their function is not required in tumor cells (e.g. they’re only required during development, or only in some tissues). Analyzing the frequency with which two members of a paralog family are lost would provide more direct insight into the contribution of paralog redundancy, but due to the overall rarity of passenger gene HDs, we cannot make a comprehensive assessment of co-deletions here – e.g. among paralog pairs where both genes are non-drivers, and not on the same chromosome, only two pairs are co-deleted in at least one TCGA sample. Larger cohorts would also allow us to search for patterns of mutual exclusivity of HDs to identify genetic interactions – this has been applied for identifying interactions between driver genes (Ciriello *et al*, 2012; Canisius *et al*, 2016), but is more challenging for interactions between non-driver genes, which are much less frequently altered.

Recent work has demonstrated that synthetic lethal interactions between paralog pairs in cancer cell lines are relatively common (Dede *et al*, 2020; Thompson *et al*, 2021; De Kegel *et al*, 2021; Parrish *et al*, 2021). This can potentially be exploited to develop targeted therapies that selectively kill tumor cells with recurrently deleted paralogs (Oike *et al*, 2014; Muller *et al*, 2012; Lord *et al*, 2020; D’Antonio *et al*, 2013). Our results provide an explanation for why the loss of paralogs is so common in tumor genomes and hence why such approaches may be broadly applicable.

One potential limitation of our analysis is that some of the driver and non-driver genes may be misclassified. For instance, it is likely that there are some frequently deleted TSGs in the patient cohorts that are not currently included in the Cancer Gene Census or identified by (Bailey *et al*, 2018) – although the latter analyzed the TCGA cohort specifically, TSGs were mainly identified from mutation rather than copy number data. However, given that we analyzed all non-driver genes collectively, it is unlikely that our conclusions would be altered significantly by the addition or removal of a small number of driver annotations. A second limitation, indicated by our saturation analysis (Fig. S3), is that there are likely many gene HDs that are unobserved due to limited sample size, rather than negative selection. As larger patient cohorts are assembled it will become possible to dissect further the influences of paralogy and other factors – e.g. functional groups that cannot be lost – on HD frequency.

## Methods

### Identifying HDs in TCGA tumor samples

We use 9,966 tumor samples from the Cancer Genome Atlas (Cancer Genome Atlas Research Network *et al*, 2013) that have been processed with ASCAT (Van Loo *et al*, 2010); downloaded from https://github.com/VanLoo-lab/ascat/tree/master/ReleasedData/TCGA_SNP6_hg19. The ASCAT results were filtered to those for which QC==‘Pass’. We straightforwardly identified HD segments as those where both the major and minor allele copy number is 0. To map genes to HD segments we used the annotations from the consensus coding sequence database (GRCh37.p13 assembly, CCDS release 15) (Pujar *et al*, 2018) and considered a gene to be deleted if its coding region is fully within the bounds of an HD segment. We consider a gene’s coding region to range from the start of the first exon to the end of the last exon of its longest transcript. Six genes are fully or partially outside the genomic region mapped by the SNP6 microarray, so we exclude these from analysis. We dropped 15 ‘hyper-deleted’ samples with over 100 homozygously deleted genes from our analysis as this far exceeds the number of deleted genes in the majority of samples (Fig. S1A).

### Identifying HDs in ICGC tumor samples

We use the 1,782 white-listed ICGC samples from the PCAWG study (ICGC/TCGA Pan-Cancer Analysis of Whole Genomes Consortium, 2020). Although the PCAWG study included both ICGC and TCGA samples, we only use the ICGC samples to ensure that this constitutes an independent dataset. The segment-level allele-specific copy number calls in PCAWG are based on the consensus of six algorithms applied to whole genome sequencing data. We then followed the same procedure as above for identifying homozygous deletions, and in this case dropped 8 ‘hyper-deleted’ samples with over 100 deleted genes – these again represent extreme outlier samples (Fig. S1B).

### Driver genes

We compiled a broad list of 652 driver genes from the Cancer Gene Census (CGC) (Sondka *et al*, 2018) and Bailey et al. (Bailey *et al*, 2018), including both Tier 1 and 2 entries from the CGC. We considered genes to be potential TSGs if either: (1) the ‘Role in Cancer’ field in the CGC dataset included ‘TSG’ or was blank; or (2) the gene was labeled as ‘tsg’ or ‘likely tsg’, or was ‘rescued’, in the Bailey et al. dataset. This gave us a list of 411 TSGs from which we identified (recurrently) deleted TSGs for the analyses related to Fig. 3 and 4.

### Identifying and annotating passenger genes

We started with a comprehensive list of autosomal protein-coding genes from the HGNC (Braschi *et al*, 2018) (2021-07-01 release), and filtered this down to genes for which there are annotations in the hg19 version of the CCDS. From this list we then filtered out the driver genes described above as well as 16 genes whose coding regions, according to CCDS, at least partially overlap the coding region of one of those driver genes. A gene was annotated as a paralog if it has at least one paralog in Ensembl 93 with which it shares at least 20% reciprocal sequence identity (Zerbino *et al*, 2018). Maximum sequence identity for each paralog gene was also obtained from Ensembl 93. We labeled paralog pairs as whole genome duplicates (WGDs) based on their inclusion in the list of WGDs identified by (Makino & McLysaght, 2010) or the strict list of WGDs in the OHNOLOGS v2 resource (Singh & Isambert, 2020). All genes that are part of at least one such WGD pair are labeled as WGD, while the rest are labeled as SSD.

### Identifying centromere and telomere-bound HDs in the TCGA cohort

As the copy number profiles for the TCGA data are computed from SNP6 array data, the genomic regions for which copy number is available are limited by the coverage of the array probes. Thus, to identify likely centromere- and telomere-bound HDs we identified the boundaries, in terms of genomic coordinates, of the data output by ASCAT. We consider a segment to be telomere-bound if it starts or ends at the outer boundaries, i.e. the first or last seen genomic coordinates respectively. Similarly, we consider a segment to be centromere-bound if it ends/starts at the maximum/minimum observed coordinate before/after the centromere assembly gap in hg19/GRCh37. Coordinates of telomere and centromere assembly gaps were obtained from the UCSC Genome Browser.

### Logistic regression models for 1+ and 3+ passenger HDs

For all passenger genes we fit logistic regression models with this form:

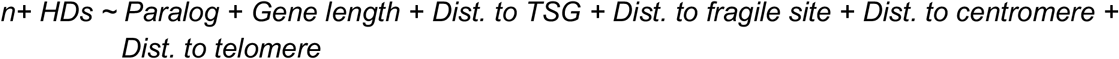

Where the model variables are as follows:

- *n+ HDs* is a Boolean variable describing whether a gene has 1+ or 3+ (vs. 0) HDs across both cohorts (TCGA and ICGC).
- *Paralog* is a Boolean variable describing whether the gene has at least one paralog.
- *Gene length* is the number of bases from the start of the first exon to the end of the last exon for the longest transcript associated with the gene (z-scored).
- *Dist. to TSG* is the inverted distance from the gene to the nearest recurrently deleted (i.e. 3+ HDs) TSG (z-scored). We cap the distance at which a TSG could influence passenger deletion at 10Mb, as this corresponds to the ~99th percentile of observed HD segment lengths across both cohorts, and then use 10 minus distance as the variable’s value. Thus a value of 10 implies the gene is right next to a recurrently deleted TSG, and a value of 0 implies the gene is >10Mb from a recurrently deleted TSG, or on a chromosome without a recurrently deleted TSG.
- *Dist. to fragile site* is the inverted distance from the gene to the nearest fragile site (z-scored). Distance is computed as for *Dist. to TSG*.
- *Dist. to telomere* and *Dist. to centromere* are the inverted distances from the gene to the nearest telomere and centromere respectively (z-scored). These distances are also computed as for *Dist. to TSG*.

For paralog passenger genes only, we fit logistic regression models with this form:

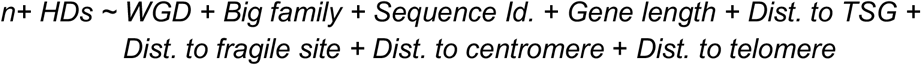

Where the new model variables (not described above) are as follows:

- *WGD* is a Boolean variable that indicates whether the gene is part of a paralog pair that originated with a whole genome duplication.
- *Big family* is a Boolean variable that indicates whether the gene has at least 4 paralogs (i.e. has a family size greater than the median of 3) with which it shares at least 20% reciprocal sequence identity.
- *Sequence Id*. is the sequence identity the gene shares with its closest paralog, i.e. its most sequence identical paralog (z-scored).

To fit these models we used the ‘Logit’ function from the python statsmodels package (version 0.11.0) (Seabold & Perktold, 2010) with default parameters.

### Identifying essential genes

To identify broadly essential genes from the DepMap CRISPR screens we used the gene dependency scores from (Table S2, (De Kegel *et al*, 2021)). These gene dependency scores were computed with CERES (Meyers *et al*, 2017) after filtering out potential multi-targeting single guide RNAs, which disproportionately affect paralog dependency scores. Broadly essential genes were identified as those with CERES score < −0.6 in at least 90% of cell lines (n=969). Essential genes from the Blomen et al. study (n=1,664) are those that were identified as essential in both the KBM7 and HAP1 cell lines (Table S3 (Blomen *et al*, 2015)). Finally essential genes from the Han et al. study (n=1,429) are those that fall within the top 2000 (based on T-score) significant negative hits in at least 2 out of the 3 cell lines screened (Han *et al*, 2020). The numbers of genes indicated refer to the number of (autosomal) essential genes after merging with our full gene list.

## Supporting information

Table S1. Binary matrix of gene HD for TCGA samples.

Table S2. Binary matrix of gene HD for ICGC samples.

Table S3. Annotated passenger genes.

## Data and Code Access

Notebooks with the code for all figures and statistical analysis are available at: https://github.com/cancergenetics/paralog_HDs

## Acknowledgements

We thank the Peter van Loo lab for making the ASCAT v3 profiles of the TCGA samples publicly available, and Dr. Tom Lesluyes in particular for answering questions regarding this data. We thank Dr. Rory Johnson, Dr. David Adams, Prof. Kenneth Wolfe, and members of the Ryan lab for providing helpful feedback on the manuscript. BDK and CJR were funded through an Irish Research Council 2017/2018 Laureate Award awarded to CJR. CJR is also supported by Science Foundation Ireland under grant number 20/FFP-P/8641.

## Author’s contributions

BDK and CJR developed the hypothesis and methodology. BDK performed data curation and analysis as well as data visualization under the supervision of CJR. BDK and CJR drafted the manuscript; all author(s) read and approved the final manuscript.

## Supplemental Figures

**Fig S1.**
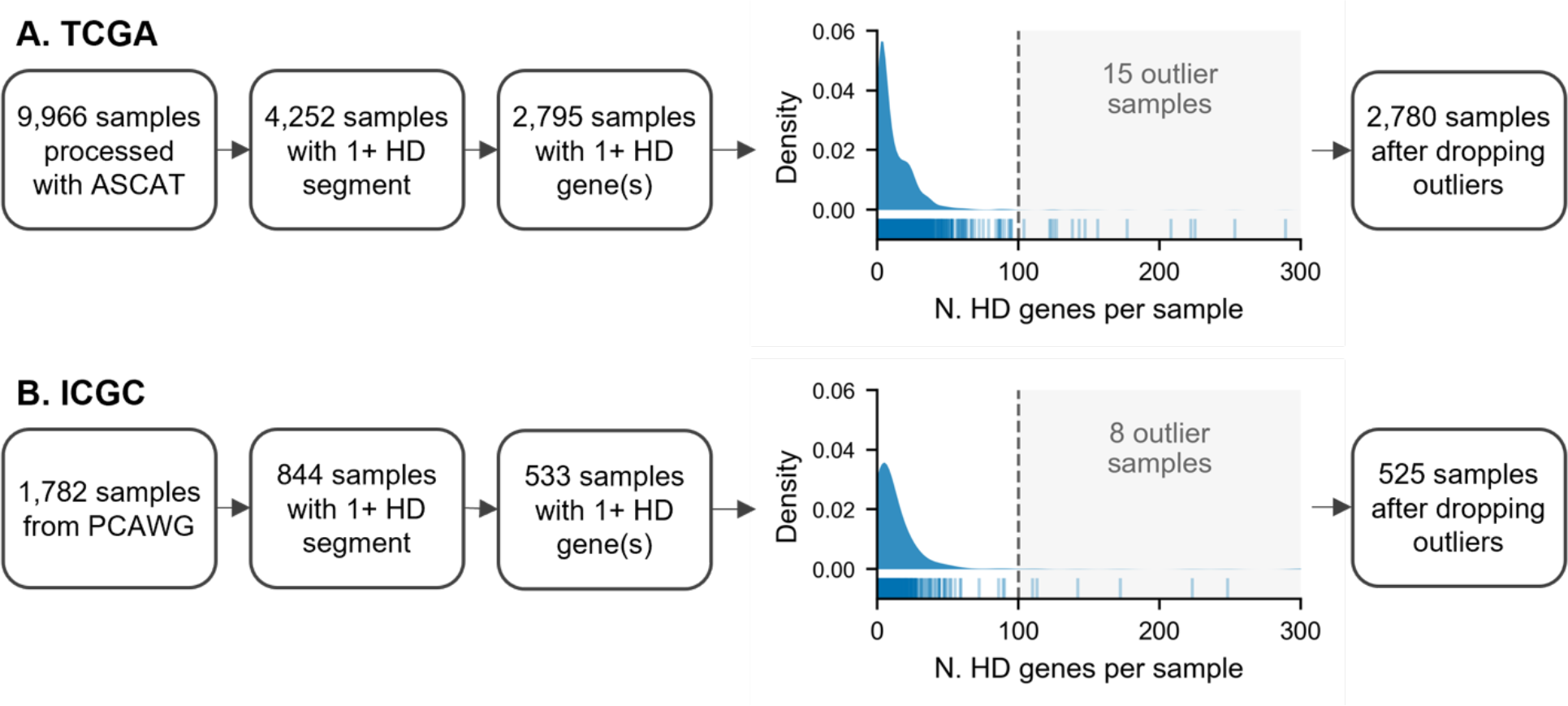
Workflow for tumor samples used in this study. (A) Flowchart for the number of TCGA tumor samples used for HD analysis. The density/tick plot shows the distribution of the number of HD genes per sample; only samples with at least 1 gene HD are shown in this plot. Samples to the right of the dotted line were marked as outliers and dropped from further analysis. (B) Same as (A) but for ICGC tumor samples from the PCAWG study.

**Fig S2.**
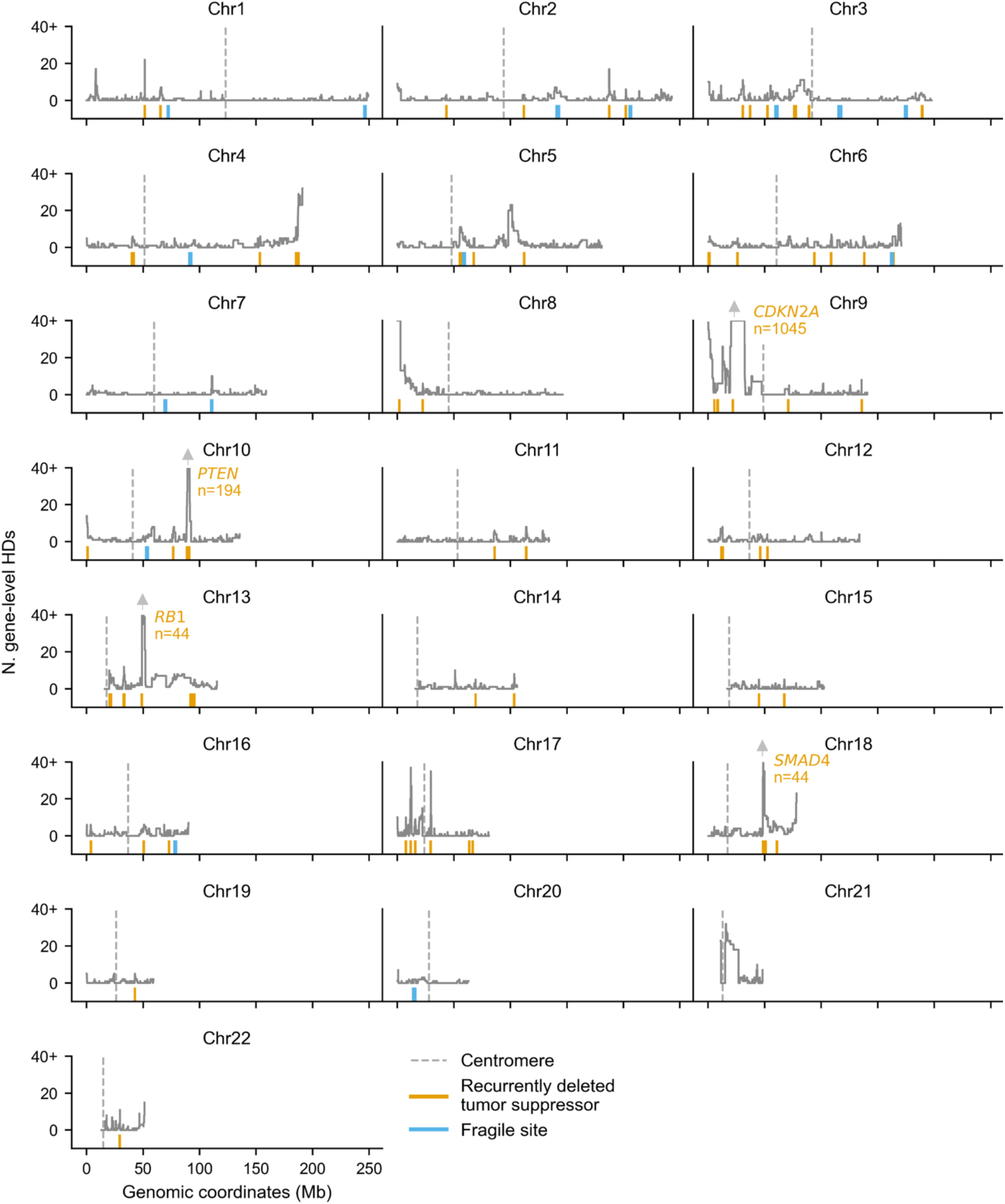
Gene-level homozygous deletion frequency across the genome. Line plots showing, for each gene plotted according to its genomic location, the number of TCGA tumor samples in which the gene is fully homozygously deleted. The number of samples is capped at 40 for visualization purposes. Orange ticks show the location of all TSGs with at least 3 HDs; four TSG peaks are annotated with the gene symbol and number of HDs. Fragile sites are denoted by blue ticks and centromeres by dotted gray lines.

**Fig S3.**
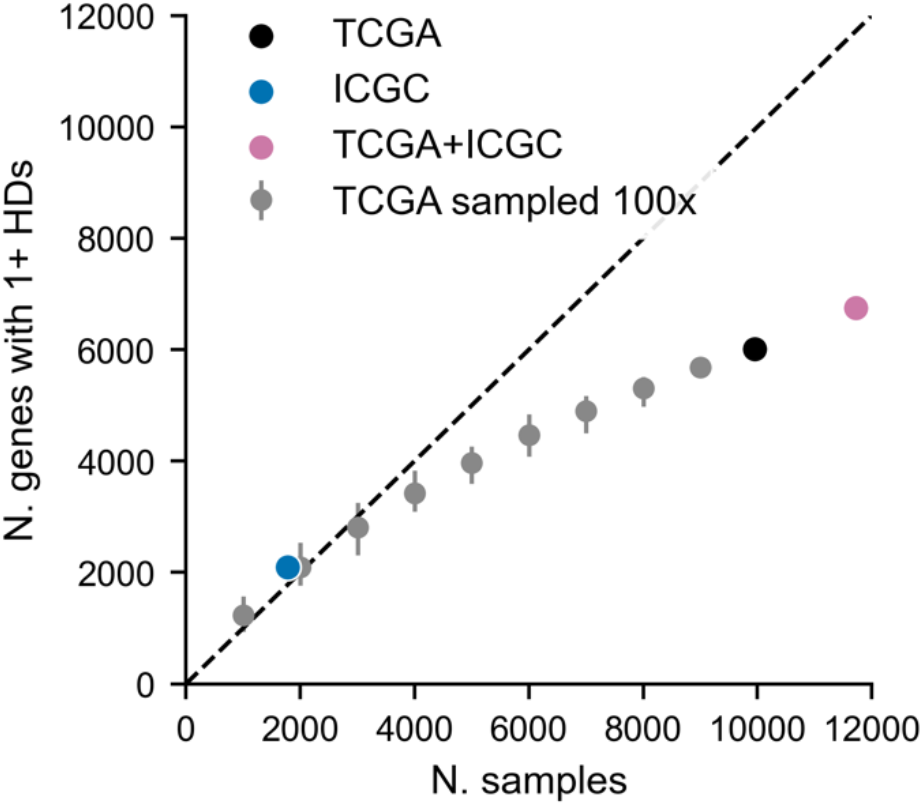
Gene HD saturation analysis. Dot plot showing the number of unique gene HDs observed (y-axis) for increasing numbers of tumor samples (x-axis). The black dot shows the actual number of unique gene HDs observed in the TCGA cohort. Gray dots are the result of down-sampling the TCGA cohort and error bars indicate the minimum and maximum values observed from 100 random samplings. The blue dot shows the actual number of unique gene HDs observed in the ICGC cohort, while the pink dot indicates the number of gene HDs that are observed when combining the TCGA and ICGC cohorts into one dataset.

**Fig S4.**
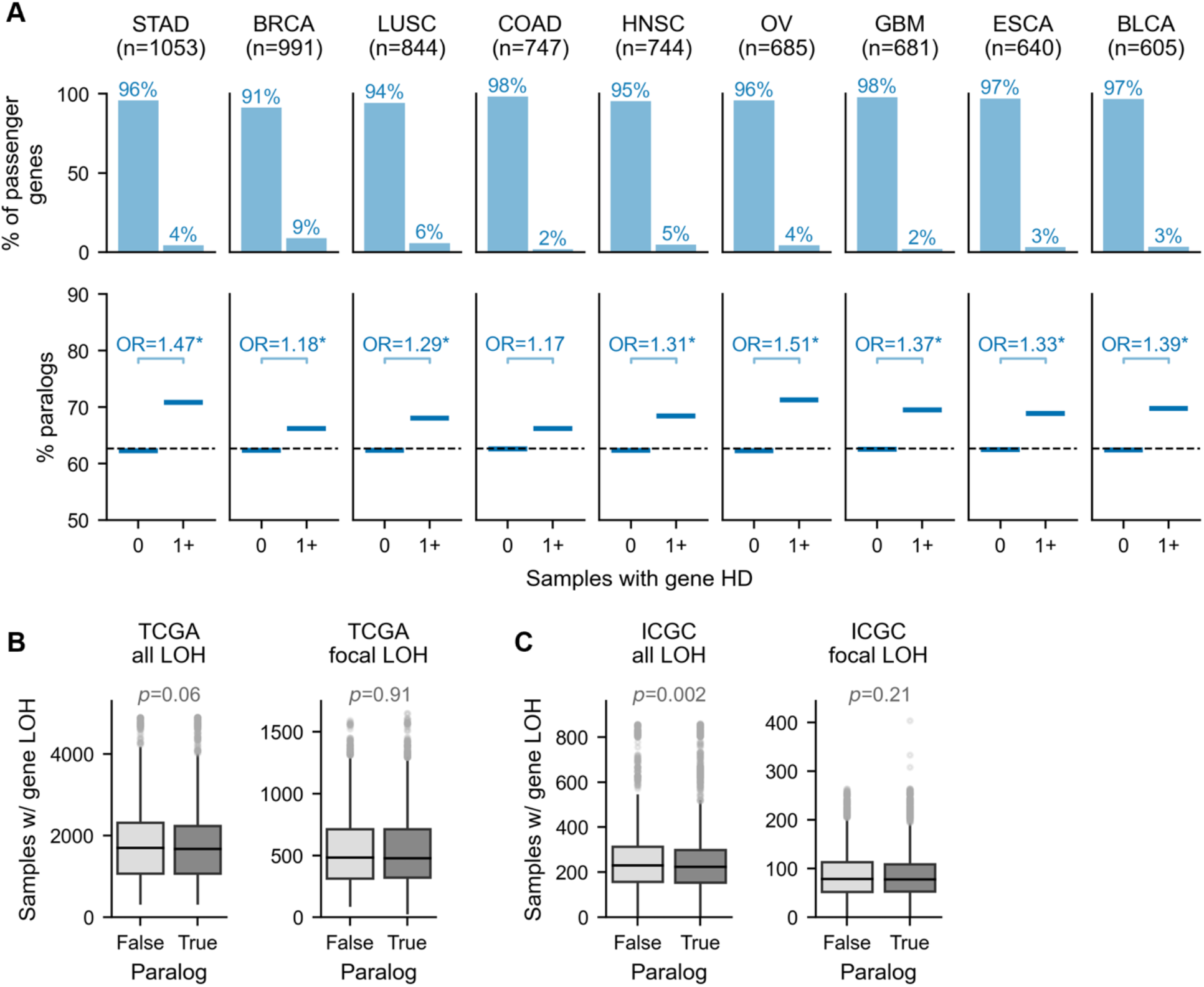
Paralog passengers are more likely to be subject to homozygous but not hemizygous deletion. (A) Same as Fig. 2A but for TCGA tumor samples stratified by cancer type. Plots are shown for all cancer types for which there are at least 600 samples. (B) Boxplots showing the number of TCGA samples in which gene-level LOH is observed for singleton vs. paralog passenger genes, when considering all LOH segments (left), or only focal LOH segments (right). The *p*-values shown are from MWU tests comparing paralogs and singletons. (C) Same as (B) but for ICGC tumor samples.

**Fig S5.**
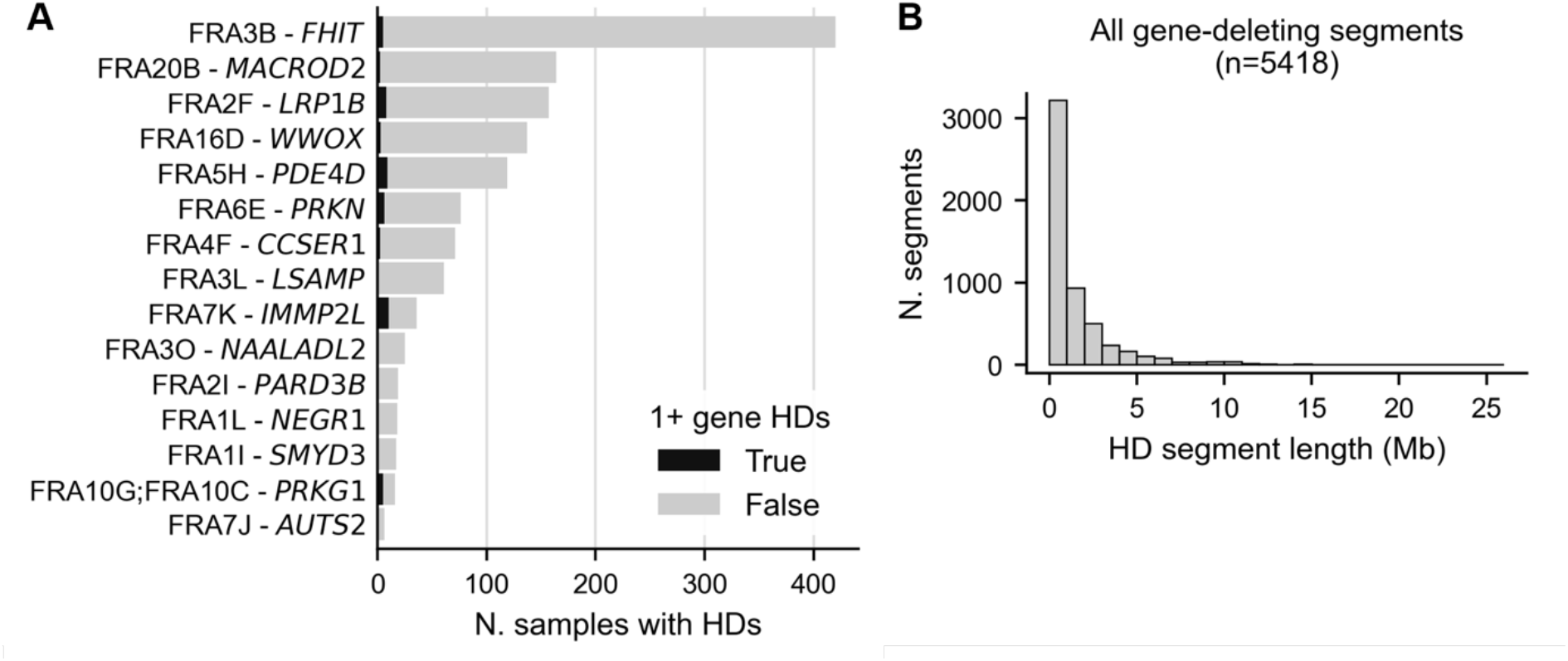
Characterization of HD segments. (A) Bar plot showing the number of TCGA samples with an HD overlapping each of 15 major fragile sites, with the section of the bar colored black indicating the number of HDs that result in at least 1 full gene HD. Fragile sites are listed with their name (e.g. FRA3B) and the longest gene they contain (e.g. *FHIT*). (B) Histogram showing the distribution of the lengths of all HD segments from both cohorts that fully delete at least 1 gene.

## Supplemental Tables

**Table S1. Binary matrix of gene HD for TCGA samples**. Columns are TCGA patient IDs (excluding outlier samples, see Fig. S1A), rows are HGNC gene symbols.

**Table S2. Binary matrix of gene HD for ICGC samples**. Columns are ICGC donor IDs (excluding outlier samples, see Fig. S1B), rows are HGNC gene symbols.

**Table S3. Annotated passenger genes**. For each passenger gene: HGNC gene symbol, Entrez ID, Ensembl ID, number of HDs across the TCGA and ICGC cohorts combined, and all features used in the logistic regression models (pre-z-scoring, see Methods).

